# Shallow sequencing (Shall-seq) procedure: A method for determining the genetic lineages and detecting antimicrobial resistance genes of antimicrobial-resistant bacteria using minimum Nanopore sequencing data

**DOI:** 10.1101/2024.06.25.600723

**Authors:** Nobuyoshi Yagi, Nanase Miyagi, Ayumi Uechi, Itaru Hirai

**Affiliations:** Laboratory of Clinical physiology, School of Health Sciences, Faculty of Medicine, University of the Ryukyus, Okinawa, Japan; Laboratory of Microbiology, School of Health Sciences, Faculty of Medicine, University of the Ryukyus, 207 Uehara, Nishihara, Okinawa 903-0215, JAPAN; Division of Clinical Laboratory and Blood Transfusion, University of the Ryukyus Hospital, 207 Uehara, Nishihara, Okinawa 903-0215, JAPAN

**Keywords:** Antimicrobial-resistant bacteria, Whole-genome sequencing, genetic characterization

## Abstract

Antimicrobial-resistant bacteria could cause nosocomial infections and outbreaks in healthcare facilities. Phylogenetic analyses based on whole-genome sequencing (WGS) could become the gold-standard method for understanding the route of antimicrobial-resistant bacterial spreading. However, generally, the WGS needs to analyze much amount of data. Therefore, sufficient resources such as budget and data analysis system are needed and it is a burden for introduction of the WGS in the routine clinical examination of pathogenic bacterial isolates. In this study, we used *Escherichia coli* as a model and evaluated whether determination of the genetic background and detection of antimicrobial-resistance genes of 29 *E. coli* clinical isolates were achieved by searching databases using sequence reads output by the Nanopore sequencer as the search keys. Consequently, only 66.2 MB data was sufficient to search for a genome sequence with ≥90% range of coverage rate. Importantly, AMR genes and plasmid replicon types were also detected with minimum data, and the detected AMR genes and phenotypes of the *E. coli* isolates did not present any discrepancy. Taken together, this shallow sequencing (Shall-seq) procedure consists of “shallow of coverage” sequencing using the Nanopore sequencer and data search using minimum data could be used to analyze bacterial isolates cost-effectively.

## Introduction

The emergence of various antimicrobial-resistant bacteria has become an important and serious public health issues of this century [1, 2]. Many previous studies have indicated the spread of antimicrobial-resistant bacteria such as extended-spectrum beta-lactamase (ESBL)-producing Enterobacterales and colistin-resistant Enterobacteriaceae in in many countries and regions [3, 4].

Nosocomial infections caused by antimicrobial-resistant bacteria and antimicrobial-resistant pathogenic bacterial outbreaks occur occasionally. To understand the distribution mechanisms of these causative bacterial isolates, genetic relatedness among the bacterial isolates was evaluated using the conventional gold-standard method, that is, pulsed-field gel electrophoresis. Bacterial whole-genome sequencing (WGS) has recently become a major method used to study the molecular epidemiology of antimicrobial-resistant bacteria. WGS can not only determine the antimicrobial-resistant bacterial species and genetic lineage, but also detect antimicrobial resistance (AMR) genes that are contained in plasmid(s) [5, 6]. The core-genome multilocus sequence typing (MLST) and core-genome single nucleotide polymorphism (SNP) analyses have been applied to study the molecular epidemiology of antimicrobial-resistant bacteria [7, 8]. Therefore, if the introduction of WGS into routine clinical examinations could enable to monitor causative bacterial isolates, which might enable rapid management of nosocomial infections and outbreaks. However, high cost is a bottleneck to utilize the WGS for a routine clinical examination. Thus, a novel labor-, power-, and cost-effective analytical method is required to address AMR.

Meanwhile, bacterial whole-genome sequences have been accumulating rapidly in public genetic databases, such as GenBank and RefSeq [9, 10]. Therefore, many genome sequence records of bacterial species commonly detected in healthcare-associated facilities, such as *E. coli* and ESKAPE, have possibly been deposited in the public databases.

We aimed to establish a novel labor-, power-, and cost-effective method for analyzing antimicrobial-resistant bacteria. Accordingly, we determined the minimum amount of sequence data required to select bacterial genome sequences from the public database and detect AMR gene sequences in the bacterial isolate genomes. In this study, we propose a shallow sequencing (Shall-seq) procedure that determines bacterial genetic lineages and detects the AMR genes harbored by antimicrobial-resistant bacteria.

## Materials and Methods

### Bacterial isolates

A total of 29 cefotaxime-resistant *E. coli* isolates obtained from two hospitals in Okinawa Prefecture, Japan, between 2013 and 2023 were used in this study.

### *E. coli* reference genome sequences

The complete *E. coli* genome sequences were retrieved from nucleotide database, “RefSeq” [10], using the keyword “complete sequence,” and filtered by taxonomy, “*Escherichia coli*.” Short sequences such as less than 4,000,000 bp and inaccurate sequences containing “N” nucleotide were removed, and the remaining 3,186 *E. coli* genome sequences were used as the reference genome sequences (**Table S1**).

Phylogenetic groups and sequence types (STs) of the 3,186 *E. coli* genome sequences were determined using EzClermont [11] and Staramr [12] with the MLST [13] database, respectively (**Table S1**).

### Library preparation for the Nanopore sequencing

The whole DNA of the *E. coli* isolates was purified using a Monarch DNA purification kit (New England Biolabs Japan Inc., Tokyo, Japan). DNA concentrations were measured using Qubit™ 4 fluorometer with Qubit™ 1× dsDNA High Sensitivity (HS) kit (Thermo Fisher Scientific K.K., Tokyo, Japan). The library was prepared using 250 ng extracted DNA with the Ligation Sequencing Kit SQK-LSK109 (Oxford Nanopore Technologies, Oxford, UK) and the native barcoding expansion kit EXP-NBD104 (Oxford Nanopore Technologies) according to the manufacturer’s protocol. The library containing eight barcode-ligated DNA was loaded onto an R9.4.1 Flongle flow cell (Oxford Nanopore Technologies).

### Reference genome mapping

Sequence reads were aligned with 3,186 *E. coli* genome sequences using Minimap2 [14]. The coverage range was calculated using SAMtools with a coverage command [15], as listed in **Table S2**. The assembled genome sequence of *the E. coli* isolate was constructed using Unicycler [16].

### Collection of AMR genes and plasmid replicons

To detect AMR genes and plasmid replicon types from sequence reads, AMR genes and plasmid replicon types were collected from the AMRFinderPlus [17] and PlasmidFinder [18] nucleotide databases, respectively. Two or more >90% homologous sequences were clustered into one sequence using VSEARCH [19].

### Construction of phylogenetic tree by core-genome analysis

Among the 3,186 reference genome sequences, 260 were selected based on the phylogenetic group and ST. After adding **NZ_CP051719.1** and the draft N75 genome sequence, we phylogenetically analyzed these 260 genomes based on the core-genome using Roary [20]. The phylogenetic tree was drawn using R and RStudio with “ggtree” package.

### Shall-seq procedure

1. **Shall-seq:** The libraries for Nanopore sequencing were prepared using 250 ng DNA. Up to 16 samples could be analyzed simultaneously using a single Flongle flow cell. A total of 250 ng (approximately 16–30 ng per bacterial isolate) of mixed DNA samples from up to 16 bacterial isolates were used for library preparation.
2. **Indicator genome sequence (IGS) selection:** The resulting Nanopore sequence reads with >9 quality score and >500 bp read length were used to select an IGS. The selected sequence reads obtained from each bacterial isolate were aligned with each *E. coli* genome sequence, as described above. In this study, the range of coverage was calculated using SAMtools, and the genome sequence showing the highest range of coverage rate, which should be >90%, was selected as an IGS representing the phylogenetic group and ST.
3. **AMR gene and plasmid replicon type detection:** The obtained sequence reads were aligned with sequences from the collection of AMR genes and plasmid replicons using KMA [21]. The sequences of AMR genes and plasmid replicon types with ≥60% “Template coverage” and ≥90% “Template identity” were counted.

## Result and Discussion

### Evaluating the data amount required to determine the genetic lineage of antimicrobial-resistant bacteria

To determine the minimum data required to identify the most identical *E. coli* genome sequence from the 3,186 *E. coli* reference genome sequences, we performed whole-genome Nanopore sequencing of *E. coli* clinical isolate N75. The entire sequence read output (962.8 MB, equivalent to 471.4 M bp) from a single Flongle flow cell were divided 1/2 to 1/512 with the seqkit [22] using command listed in **Table S2**. The divided sequence reads were aligned with each *E. coli* genome sequence. The range of coverage rates decreased depending on the sequence data amount (**Fig. 1**). However, the range of coverage rate for 66.4 MB (32.6 M bp) was not substantially different from that for >66.4 MB sequence data. Moreover, the *E. coli* isolate N75 sequence data selected **NZ_CP051719.1** from the 3,186 *E. coli* reference genome sequences, but even if only 8.5 MB (4.2 Mbp) sequence data were used for the alignment, the same **NZ_CP051719.1** was selected.

**Fig. 1.**
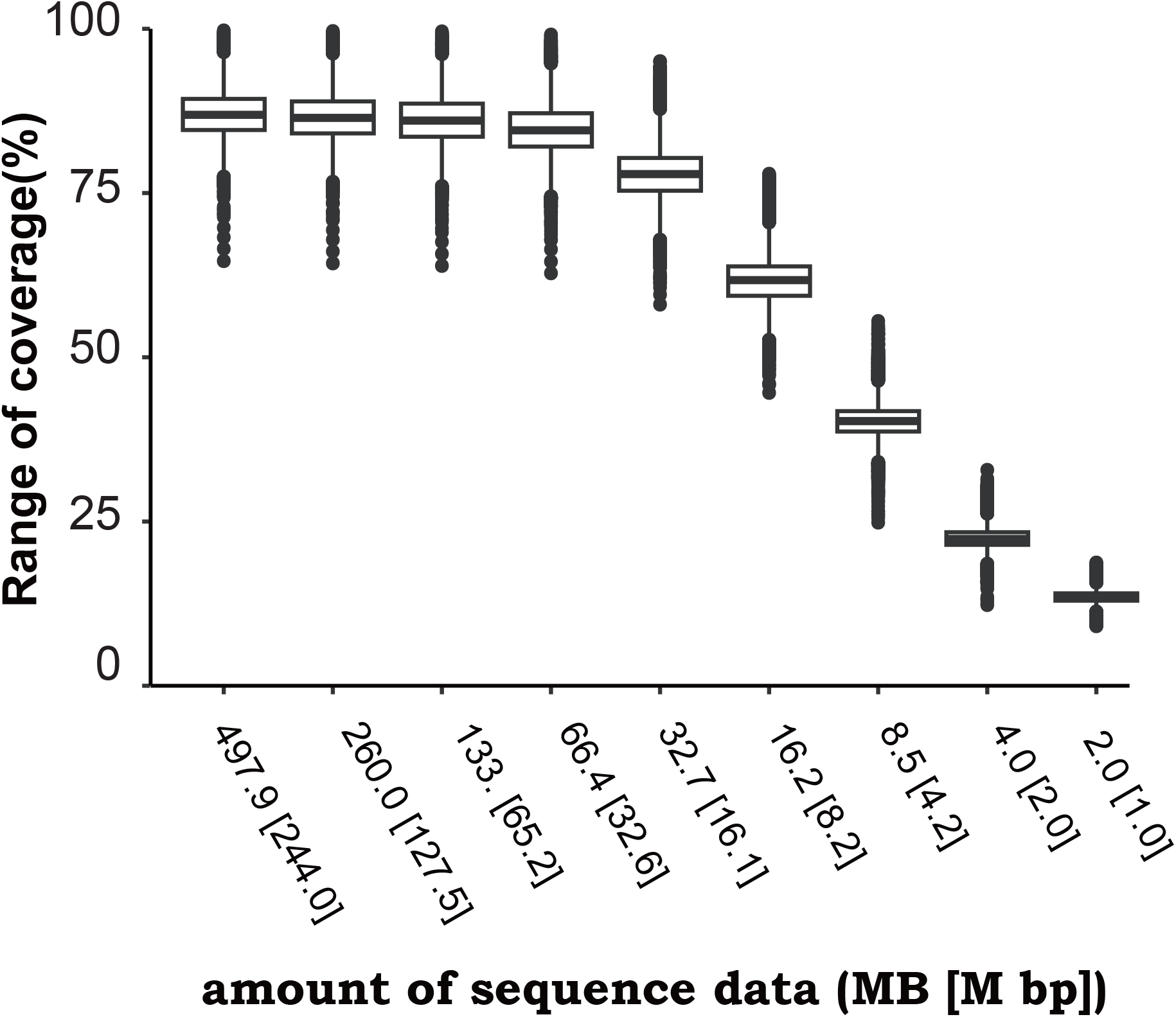
The effect of analysis using less sequence data. Boxplots show the impact of sequence data reduction on the coverage range analyzed by reference mapping against 3,186 *Escherichia coli* genomes individually. X-axis represents the sequence data used for alignment with 3,186 reference genome sequences, and y-axis represents the coverage range against *E. coli* reference genome sequences.

We then evaluated the location of **NZ_CP051719.1** and the assembled *E. coli* isolate N75 genome sequence in the phylogenetic tree constructed using the 260 *E. coli* genome sequences. **NZ_CP051719.1** and the assembled *E. coli* isolate N75 genome sequence were in very close branches of the phylogenetic group B2 (**Fig. 2**). Consistently, BLAST indicated a cover % of between **NZ_CP051719.1** and the assembled genome sequence of the *E. coli* isolate N75 was more than 99%.

**Fig. 2.**
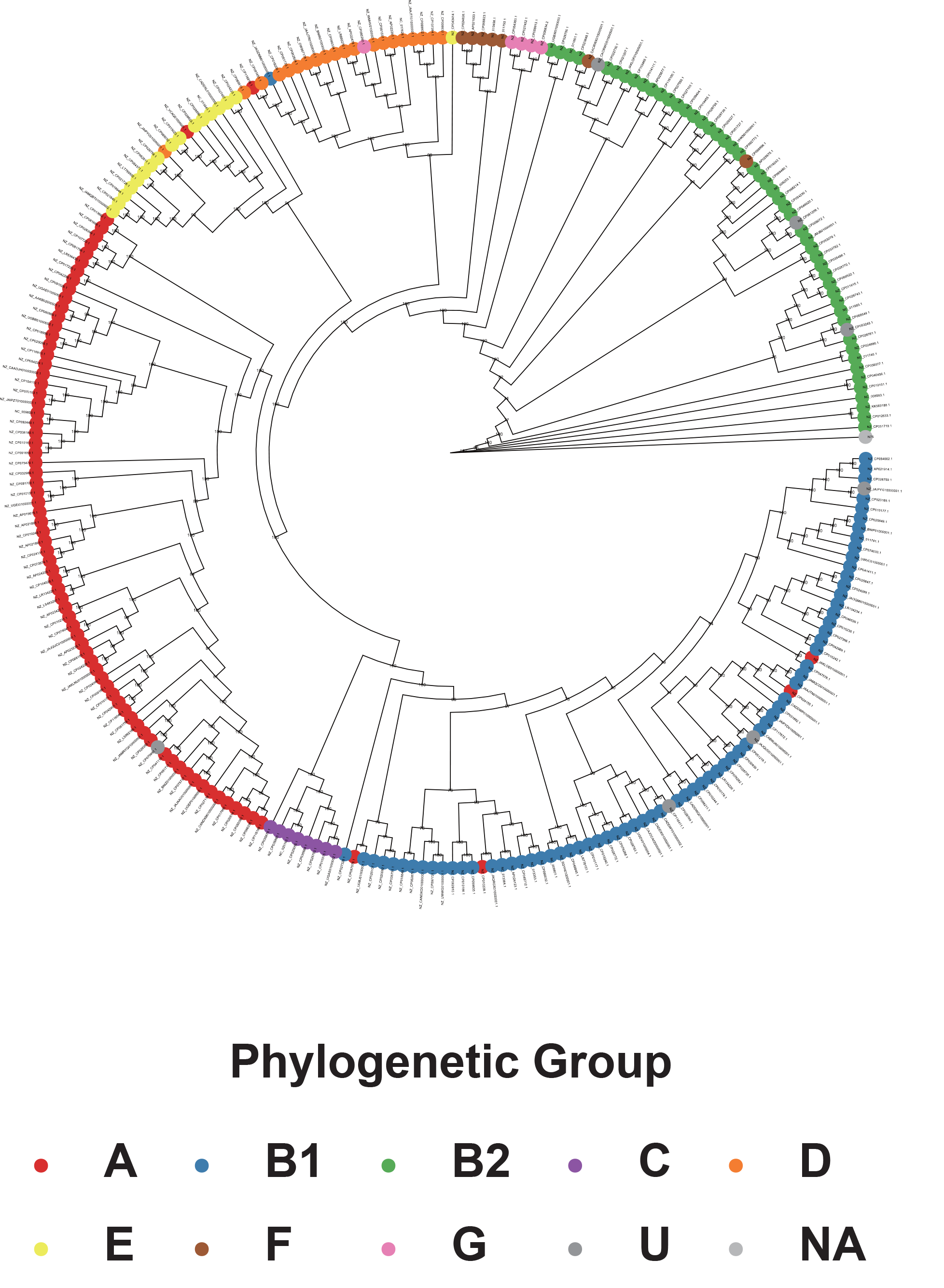
The phylogenetic tree constructed by core-genome analysis. To compare the genome sequences of **NZ_CP051719.1** and **N75**, the phylogenetic tree having 262 *Escherichia coli* genome sequences including **NZ_CP051719.1**, the assembled genome sequence of the *E. coli* clinical isolate **N75**, and 260 reference genome sequences selected based on the differences of their phylogenetic group and ST among the 3,186 *E. coli* reference genome sequences was constructed using Roary.

These results suggested that 66.4 MB (32.6 M bp) was the minimum data amount required to identify reference genome sequences homologous to clinical isolate genome sequences from the database.

### Examination of the 29 antimicrobial-resistant *E. coli* clinical isolates by the Shall-seq procedure

Our results suggest that 16 samples can be analyzed simultaneously using a single Flongle flow cell. However, we simultaneously analyzed eight samples in this study, considering the fluctuations in the amount of sequence data obtained.

For further evaluation, we analyzed total of 29 *E. coli* clinical isolates obtained from hospitals in Okinawa Prefecture, Japan. The obtained sequence reads were used to select IGSs and detect the *AMR* gene and plasmid replicon types (**Table 1**). Consequently, 11 IGSs were selected for this study (**Table 1**). Among them, 11 (81.8%), 1 (9.1%), and 1 (9.1%) IGSs were classified into phylogenetic groups B2, D, and F, respectively. A total of 174 AMR genes were detected in the 29 *E. coli* clinical isolates. On average, six *AMR* genes were detected in each antimicrobial-resistant *E. coli* clinical isolate, and *mph*(A) was the most prevalent gene (**Table 1**). Importantly, the phenotypes of the 29 antimicrobial-resistant *E. coli* isolates and *AMR* genes detected in their sequence data did not have major discrepancies.

**Table 1.**
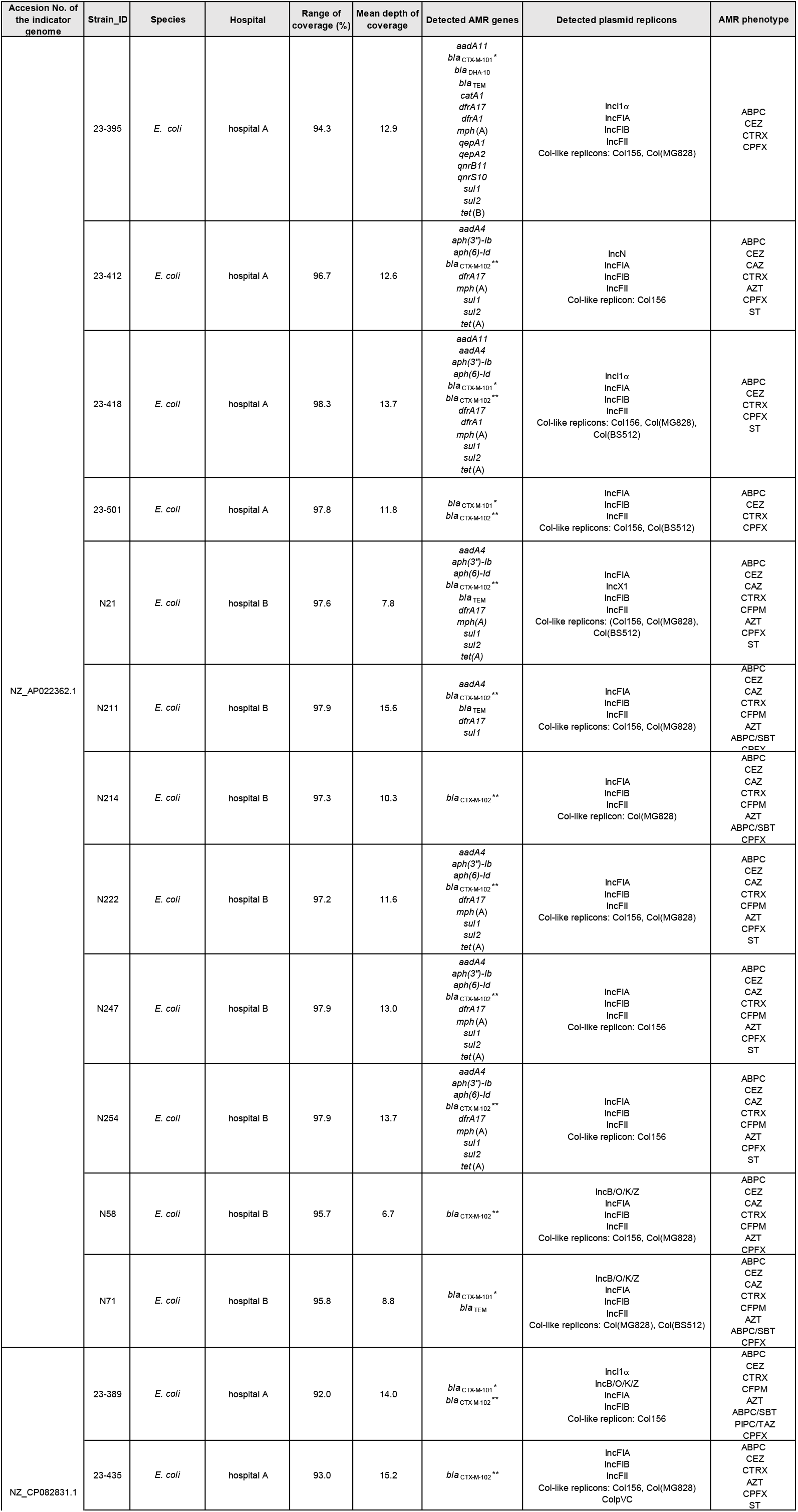
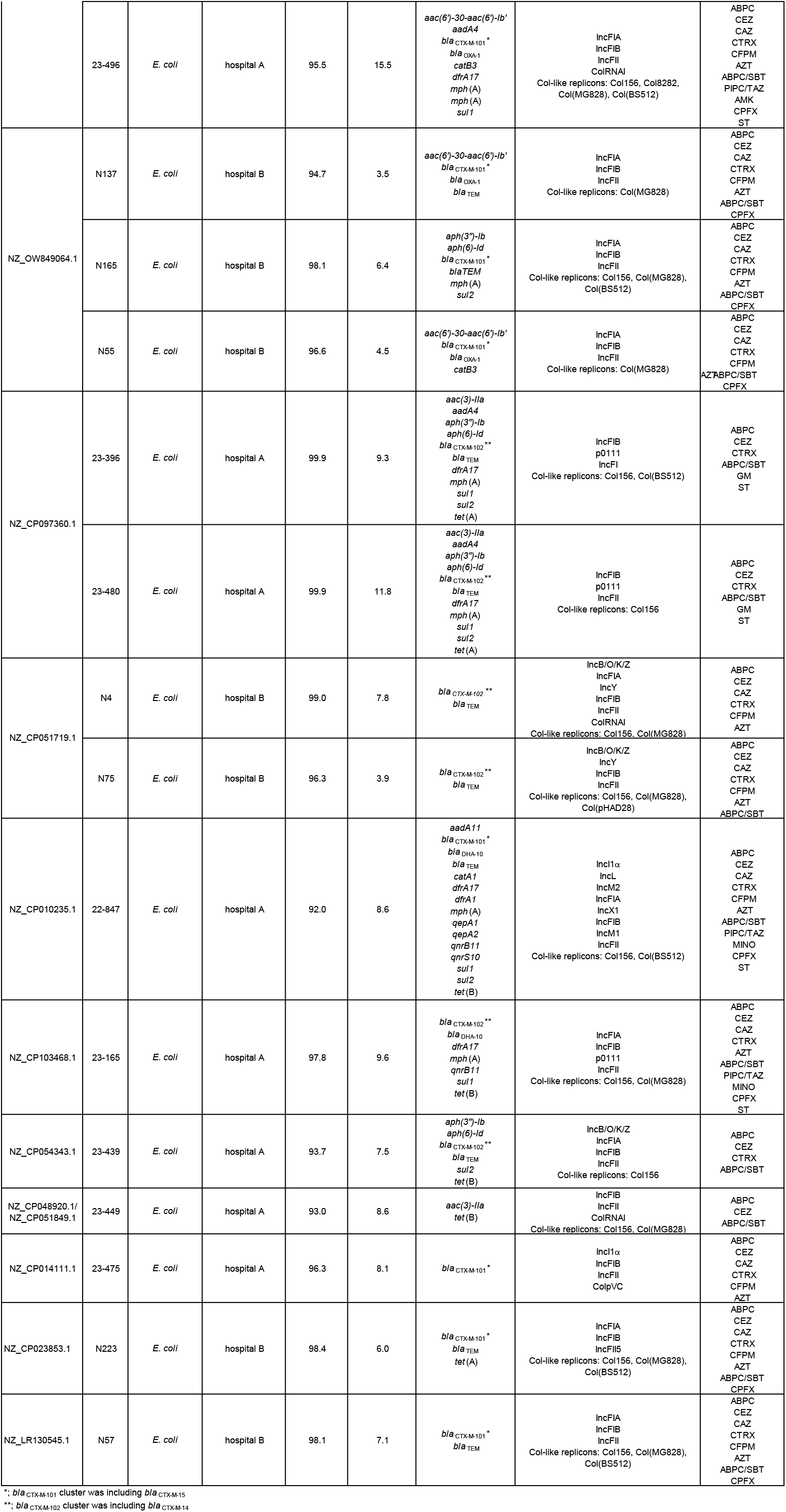
the imformation of E. coli isolates analyzed in this study.

Interestingly, **NZ_AP022362.1, NZ_LT632320.1**, and **NZ_CP082831.1**, were selected based on sequence data obtained from 11 (37.9%), 4 (13.8%), and 3 (10.3%) of the 29 examined *E. coli* clinical isolates, respectively. Since the number and species of the detected *AMR* genes in these clinical isolates were not completely matched, it is not highly possible that the three IGSs were selected due to antimicrobial-resistant bacterial contamination and sample mixing. This means that the Shall-seq procedure can classify antimicrobial-resistant bacterial isolates, at least at the ST level.

## Conclusion

The selection of the IGS homologous to clinical isolates from public nucleotide database using the Shall-seq procedure could be a cost-effective WGS, which might facilitate introducing WGS as a routine clinical examination. Consequently, it might facilitate rapid managements and interventions in the case of nosocomial infections and outbreaks caused by antimicrobial-resistant bacteria.

Although a high error rate is considered a disadvantage of Nanopore sequencing, its accuracy has recently improved [23, 24]. This improved accuracy encourages using the Shall-seq procedure to monitor antimicrobial-resistant bacteria in niches other than healthcare-associated facilities, such as communities and natural environments.

## Data availability

The sequence reads used in this study are available under BioProject accession number **PRJDB17430**.

